# An epigenetic clock to estimate the age of living beluga whales

**DOI:** 10.1101/2020.09.28.317610

**Authors:** Eleanor K. Bors, C. Scott Baker, Paul R. Wade, Kaimyn O’Neill, Kim E.W. Shelden, Michael J. Thompson, Zhe Fei, Simon Jarman, Steve Horvath

**Affiliations:** Marine Mammal Institute, Oregon State University, Newport, OR, USA, 97365; Marine Mammal Laboratory, Alaska Fisheries Science Center, National Marine Fisheries Service, National Oceanographic and Atmospheric Administration, Seattle, WA, USA; Molecular, Cell and Developmental Biology, University of California Los Angeles, Los Angeles, CA 90095, USA; Department of Biostatistics, School of Public Health, University of California-Los Angeles, Los Angeles, CA, USA; School of Biological Sciences, University of Western Australia, Perth, Australia; Department of Human Genetics, Gonda Research Center, David Geffen School of Medicine, Los Angeles, CA, USA

**Keywords:** DNA methylation, cetaceans, aging, endangered species, Cook Inlet, Alaska, conservation

## Abstract

**Introduction:** DNA methylation data facilitate the development of accurate molecular estimators of chronological age, or ‘epigenetic clocks.’ We present a robust epigenetic clock for the beluga whale, *Delphinapterus leucas,* developed for an endangered population in Cook Inlet, Alaska, USA.

**Methods and Results:** We used a custom methylation array to measure methylation levels at 37,491 cytosine-guanine sites (CpGs) from skin samples of dead whales (n = 67) whose chronological ages were estimated based on tooth growth layer groups. Using these calibration data, a penalized regression model selected 23 CpGs, providing an R^2^ = 0.92 for the training data; and an R^2^ = 0.74 and median absolute age error = 2.9 years for the leave one out cross-validation. We applied the epigenetic clock to an independent data set of 38 skin samples collected with a biopsy dart from living whales between 2016 and 2018. Age estimates ranged from 11 to 27 years. We also report sex correlations in CpG data and describe an approach of identifying the sex of an animal using DNA methylation.

**Discussion:** The epigenetic estimators of age and sex presented here have broad applications for conservation and management of Cook Inlet beluga whales and potentially other cetaceans.

## INTRODUCTION

Age is a fundamental life-history parameter in organismal biology, population dynamics, and ecology. The age of an animal is important for understanding characteristics such as age of reproductive maturity, fecundity rates, and survival rates. These characteristics can vary between healthy and compromised populations. Moreover, knowing the age of animals in a population can improve the study of population dynamics. For example, age-specific estimates of fecundity and survival can be used to predict population growth rate (Brault and Caswell, 1993), and the age structure of a population can imply past trajectory of the population (Venuto et al., 2020). Additionally, age can enhance the interpretation of genetic analyses in some cases (*e.g.,* kinship analysis). Therefore, the ability to determine the age of animals is an important tool in wildlife studies. In cetaceans, age is critically important but often unknown due to the difficulty of determining age in long-lived, mobile species.

The development of molecular aging biomarkers (MABs) for mammals in particular has been of interest for decades (Jarman et al., 2015), with numerous lines of inquiry into the role of molecular mechanisms in aging. Molecular aging studies in cetaceans initially focused on the relationship between telomere length and age, but that line of inquiry proved unfruitful (Dunshea et al., 2011; Jarman et al., 2015; Olsen et al., 2012). Attention has turned to other MABs such as epigenetic markers, specifically DNA methylation, which have received considerable attention in recent years (De Paoli-Iseppi et al., 2017; Horvath and Raj, 2018; Jarman et al., 2015; Jylhävä et al., 2017). Epigenetics is broadly understood as the study of any gene-regulating activity that does not involve changes to a DNA sequence and can be, but is not necessarily, heritable (Pennisi, 2001). The term encompasses myriad molecular processes ranging from chromatin state to direct chemical modification of DNA (e.g., methylation). At specific cytosine-guanine dinucleotides (CpGs), the cytosine nucleotide can be methylated to generate 5-methylcytosine, a chemical modification that affects gene expression (Field et al., 2018; Razin and Cedar, 1991). Methylation levels at some of these sites have been shown to correlate with age.

Numerous DNA methylation-based age predictors, often referred to as ‘epigenetic clocks,’ have been developed for humans (*e.g.*, Bocklandt et al., 2011; Garagnani et al., 2012; Hannum et al., 2013; Horvath, 2013; Levine et al., 2018; Lin et al., 2016; Lowe et al., 2018). Because methylation patterns are known to be tissue-specific, some epigenetic clocks use a single tissue such as blood (*e.g.,* Hannum et al., 2013) while pan-tissue clocks appear to apply to all sources of DNA except sperm (*e.g.,* Horvath, 2013). In humans, clocks have been designed using varying numbers of CpG sites, from one site (*e.g.*, an age predictor based on a CpG in the *ELOVL2* gene; Garagnani et al., 2012) to several hundred sites (*e.g.,* 353 sites; Horvath, 2013). Epigenetic clocks have also been developed for other species including the mouse (Stubbs et al., 2017; Petkovich et al., 2017; Thompson et al., 2018; Meer et al., 2018), chimpanzee (Ito et al., 2018), bat (Wright et al., 2018), canid (Ito et al., 2017; Thompson et al., 2017), humpback whale (Polanowski et al., 2014), minke whale (Tanabe et al., 2020), and bottlenose dolphin (Beal et al., 2019). Accurate age estimates can be valuable for conservation efforts and species management. For example, the use of age-structure data for harbor porpoise allowed for estimation of the maximum rate of increase for the population, leading to the conclusion that bycatch mortality in commercial fisheries had led to population decline (Moore and Read, 2008).

Our focus here is the beluga whale, *Delphinapterus leucas* (Pallas, 1776). Beluga whales inhabit the circumpolar north with southernmost populations occurring in the Saint Lawrence Estuary in Eastern Canada, the Sea of Okhotsk in Eastern Russia, and Cook Inlet in Alaska, USA. The Cook Inlet (CI) beluga whale population does not migrate, is geographically and genetically isolated (O’Corry-Crowe, et al., 1997), and is the focus of conservation and management efforts (NOAA, 2016). Estimates of abundance for this population numbered over 1,000 whales in the late 1970s and early 1990s (Shelden et al., 2015). From 1994 to 1998, abundance declined steeply from 653 to 347 whales, in part, due to unregulated hunting (Hobbs et al., 2000; Mahoney and Shelden, 2000). In 2000, hunting regulations were implemented; the CI beluga population was designated as a distinct population segment (DPS), recognizing that CI beluga whales constitute a population that is ‘discrete from other populations and significant in relation to the entire taxon’ (65 FR 38788 22 June 2000); and CI beluga whales were listed as depleted under the U.S. Marine Mammal Protection Act (65 FR 34590, 21 May 2000). Eight years later, the DPS was listed as endangered under the U.S. Endangered Species Act (ESA) (73 FR 62919, 22 October 2008). Today, there are an estimated 279 individuals in the population, and the number is declining (Wade et al., 2019). The U.S. National Oceanic and Atmospheric Administration (NOAA), released the *Cook Inlet Beluga Whale Recovery Plan* in 2016 (NMFS, 2016) pursuant to the requirements of the ESA. This plan highlights the importance of determining the population age structure of CI beluga whales in order to understand growth, reproduction, and survival rates.

Age determination of beluga whales to date has relied on data derived from tooth growth layer groups (Lockyer et al., 2007; Waugh et al., 2018), a method that is also applicable to some other toothed whale species (Hamilton and Evans, 2018; Perrin and Myrick, 1980). The tooth samples required for aging studies are typically acquired from dead animals. Efforts to develop methods that estimate the age of living animals have led to the development of length-age curves for adult belugas (Vos et al., 2019) as well as fetuses and neonates (Robeck et al., 2015).

Here, we present an epigenetic clock and a sex-predictive logistic model for beluga whales based on DNA methylation data. This study leverages long-term sampling of the CI beluga population, recent development of beluga genomic resources, advances in methylation array technology, and machine learning to develop a novel method to age living beluga whales based on DNA from skin samples. The epigenetic age estimator (epigenetic clock) presented here will aid in the management and conservation of this endangered cetacean population, as demonstrated by our application of the beluga epigenetic clock to estimate the age of living beluga whales sampled with a biopsy dart.

## Methods

### Sample collection, chronological age estimation, and DNA extraction

Skin tissue samples were collected from carcasses of beluga whales that were beach-cast, stranded dead, or taken during subsistence hunting between 1992 and 2015 in Cook Inlet, Alaska, USA (NMFS Research Permit 932-1905-00/MA-009526 through the Marine Mammal Health and Stranding Response Program). Skin samples were preserved in a salt and dimethyl sulfoxide (DMSO) solution and archived at NOAA’s Southwest Fisheries Science Center in La Jolla, California, USA. A total of 69 individuals were selected for the clock calibration dataset (Table S1), and their chronological ages were estimated by counting tooth growth layer groups (Vos et al., 2019). The final calibration dataset included 67 individuals due to inconsistent molecular sex data (see below). Teeth were analyzed by at least two readers using methods validated in Lockyer et al. (2007), and a consensus age provided by NOAA was used in this study. When individuals were represented by multiple teeth in the dataset, the oldest age estimate was used to mitigate error from tooth wear (*e.g.,* the count from the tooth with the greatest number of growth layer groups; Vos et al., 2019).

Samples of skin tissue from living CI beluga whales were collected with a biopsy dart in 2016, 2017, and 2018 (NMFS ESA/MMPA Permit #20465; McGuire et al., 2017a). Biopsy samples were frozen in the field in liquid nitrogen and later subsampled at the NOAA Alaska Fisheries Science Center in Seattle, Washington, USA. Genomic DNA was extracted from tissue samples using a standard phenol-chloroform protocol modified for small skin samples by Baker et al. (1994). Extracted DNA was treated with RNAse A (1 μL of 1 mg/mL added to samples of 100 μL for 30 minutes at room temperature) and then purified and concentrated using a DNA Clean and Concentrator-5 Kit (Zymo Research Corp., USA). The concentration of genomic DNA was measured on a QUBIT 4 fluorometer (ThermoFisher Scientific, USA).

### Molecular sex identification

A multiplexed polymerase chain reaction (PCR) was used to sex individual whales in both the calibration and biopsy datasets. The PCR primers and reaction protocol followed those in Olavarría et al. (2007), which is based on Gilson et al. (1998). This assay amplifies fragments of the male-specific SRY gene and the ZFY/ZFX genes of males and females as a control band. Sex-specific bands were visualized by agarose gel electrophoresis. In three cases, molecular sex identified tissue samples that did not correlate with sex metadata in the original records (z35345, z143907, z144309). These cases have been noted and amended in the records presented here (Table S1). Two of these individuals were removed from the dataset because the molecular sex could not be reconciled with information from necropsies. One was retained because it did not conflict with known information. Therefore, the final calibration dataset included 67 individuals.

### DNA methylation measurements

Genomic DNA aliquots were sent to the UCLA Neurosciences Genomics Core facility where they were quantified and bisulfite converted using the Zymo EZ-96 DNA Methylation-Gold Kit (Zymo, Inc., USA; Cat# D5007). When possible, a total of 250 ng of genomic DNA was used for each individual (in a few cases, lower quantities were used when DNA concentration was too low to achieve 250 ng with a maximum volume loadable of 20 μL). A custom mammalian methylation array (HorvathMammalMethylChip40) assembled with 37,491 oligonucleotide probes, each 50 nucleotides long terminating in a C-G dinucleotide, was used to determine methylation state of CpGs (Arneson, Ernst, Horvath, *in prep*). Each probe was designed to cover a certain subset of species. The particular subset of species for each probe is provided in the chip manifest file can be found at Gene Expression Omnibus (GEO) at NCBI as platform GPL28271. The clock training/calibration dataset (historic samples) was evaluated in one round of array assays (one sample per array, 24 arrays per chip) and the biopsy dataset (recent skin tissue samples) was evaluated in another. Fluorescence at the terminal nucleotide of each probe was read by an Illumina iScan machine and raw data were provided in iDAT files. Raw data were normalized using the SeSAMe pipeline (Zhou et al., 2018) resulting in a methylation estimate (beta value) corresponding to each array probe for every individual in the dataset and a detection p-value corresponding to the confidence in the normalized beta value. Beta values are derived from the ratio of the fluorescence intensity of a methylated probe for a specific CpG to the total overall probe intensity (the sum of signal from both the methylated and unmethylated probes plus a constant) (Du et al., 2010). Beta values range from zero to one with a value of zero indicating that no copies of the gene were methylated.

To identify technical outliers after SeSAMe normalization, we used unsupervised hierarchical clustering analysis based on the interarray correlation. As a consequence, data for one sample was removed from the dataset and replaced with data generated for another DNA extraction of the same tissue sample that did not exhibit anomalous clustering. Data were filtered by detection p-value as calculated in SeSAMe (Zhou et al., 2018). To test the effect of p-value filtering on downstream analyses, we evaluated analyses on data with different thresholds for the number of individuals with a passable detection p-value (*e.g.,* a detection p-value of < 0.05 in more than one individual, in over 10 individuals, in over 20, etc.). Ultimately, CpG sites that had a detection p-value < 0.05 for 10 or more individuals were considered in further clock-building analyses, resulting in the use of data from 28,875 CpG probes from the array (Table S2).

### Sex-correlated CpGs and methylation-based sex prediction

The correlation between CpG methylation levels and whale sex was evaluated using Pearson’s correlation using *NymPy* in a *Python* environment. The genomic location of sex-correlated CpGs in humans was recorded to ascertain how many of these sites are on the X and Y chromosomes, and how many represent sexual dimorphism in autosomal methylation levels. A logistic model for sex prediction was built using LASSO regression in cv.*glmnet()* (alpha = 1) for binomial parameters (numerical coding was 1 = female; 0 = male).

### Age correlation of CpG sites and epigenetic clock development through elastic net regularization

Pearson’s correlations between beta values for individual CpGs and chronological age were calculated using *NumPy* (Oliphant, 2006) and *Pandas* (McKinney, 2010). The absolute values of individual CpG correlations were ranked using a custom *Python* script.

The *glmnet* package (Friedman et al., 2010) was implemented in *R* (R core Team, 2013) to fit penalized regression models. The two main parameters used in this machine learning regularization method are lambda, which is known as the regularization parameter that sets the stringency of the penalty during regularization (high lambdas lead to stronger penalization); and alpha, which is the elastic net mixing parameter that is used to determine the blend between a ridge regression (alpha = 0) and a least absolute shrinkage and selection (LASSO) regression (alpha = 1). For all runs, the lambda used was the *lambda.min* parameter calculated by *cv.glmnet* using a 10-fold cross validation. In elastic net regularization, the alpha parameter will determine the number of sites used in the clock: ridge regression will retain all the sites and LASSO regression will retain fewer sites.

Alpha values of 0.1 through 0.9 with a 0.1 interval were evaluated through multiple runs of *cv.glmnet*() (note that an alpha value of 0.0 would have yielded a model using all 28,875 CpGs). The resulting models were then used to calculate the age of each individual in the calibration dataset. The relationship between model ages and chronological ages based on tooth growth layer groups was evaluated using linear regression in *NymPy* and *Pandas* within a *Python* environment. Age error was calculated for each individual, which was defined as the difference between the estimated chronological age from tooth growth layer groups and the age prediction resulting from the multivariate linear regression model. Regression coefficients, mean absolute age error, and median absolute age error were calculated for the dataset as a whole and for each sex independently.

To evaluate the likely accuracy of each model for estimating the age of future experimental samples, leave one out cross-validation (LOOCV) was run by executing the *cv.glmnet()* program on n-1 samples, looping through each of the 67 samples. The predicted age of the omitted sample was calculated with a model developed with the remaining 66 samples. For each LOOCV iteration, a different model was generated, but the same alpha value was used in each iteration of *cv.glmnet()* (only the samples used changed). LOOCV elastic net models were run with a lambda value of *lambda.min* calculated within *cv.glmnet()* program by running a 10-fold internal cross-validation. LOOCV were executed at numerous alpha values to better understand the effect of alpha on model performance. LOOCV models were assessed in the same manner as the full elastic net regression models, using linear regression as well as age error calculations for the full data set and for each sex independently.

### Genomic location and identity of clock CpGs

The location of each clock CpG probe in the human or mouse genomes was known from methylation array design. The genomic locations of each clock CpG and flanking sequences (200 bp in both directions) were extracted from the human or mouse genomes through the NCBI genome data viewer, leveraging the RefSeq database (O’Leary et al., 2016). The extracted mouse or human sequence was then located in the beluga genome (GenBank genome scaffold accession number: ASM228892v3) with NCBI BLAST (Johnson et al., 2008; Altschul, 1990). Annotations at each CpG site, or for the closest gene, were recorded. The relative locations of each CpG in the final epigenetic clock were assessed to identify any potentially linked sites. The same methods were also used to determine the annotation of the single CpG used in the sex-prediction model.

### Age determination of living whales using skin biopsy samples

We applied the beluga epigenetic clock model to the dataset of living beluga whales sampled by biopsy dart. Beta values generated by the custom mammalian methylation array for each of the clock CpG sites were used in the beluga epigenetic clock to calculate the epigenetic ages of sampled whales. The absolute median age error from LOOCV of the calibration dataset was used as a course approximation for the potential range of the epigenetic age estimation generated by the epigenetic clock. When appropriate and possible, age estimates generated with the beluga epigenetic clock were compared to photographs of each individual and compared to subjective color-classes used to assess age in the field (McGuire et al., 2017b; McGuire et al., 2018; unpublished data, P. Wade; Figure S1).

## Results

### Chronological ages and molecular sex of clock calibration samples

The estimates of chronological ages derived from tooth growth layer groups for the 67 clock calibration samples ranged from −1 (fetus) to 49 years with a median age of 21 years (Fig 1). Previous records of sex were confirmed by PCR amplification of sex-specific primers. After correction, the ratio of males to females in our calibration dataset was 36 to 31. The median age of males in the dataset was 20 while that of females was 22. However, the three oldest samples in the dataset are males based on molecular sex data (Fig 1).

**Figure 1:**
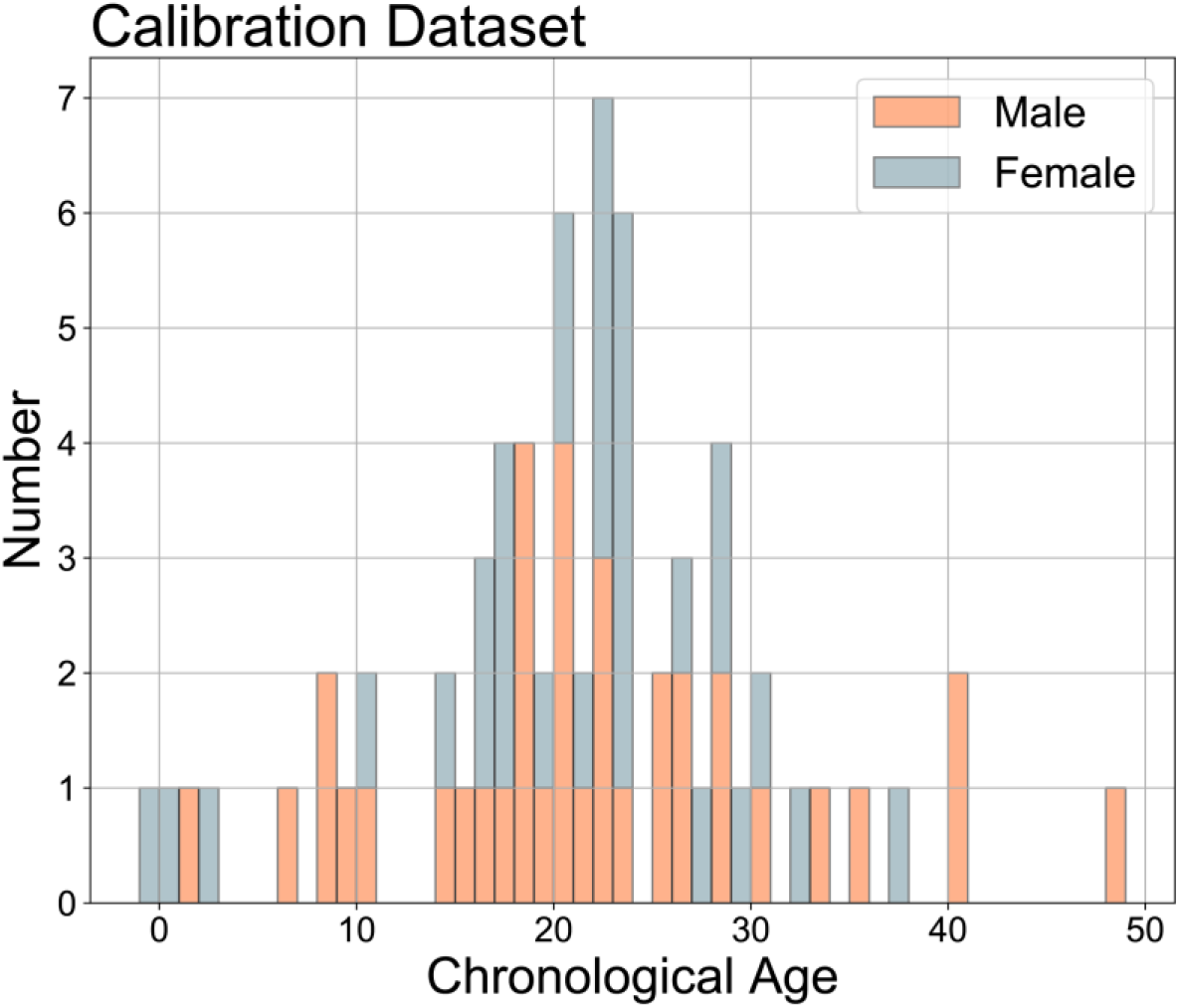
The distribution of chronological age by sex estimated from tooth growth layer groups for the calibration dataset (n=67). Note that each bin represents one year and negative ages are fetal samples.

### Sex-correlated CpG sites and methylation-based sex prediction model

In addition to correlations between methylation and age, the relationship between methylation and sex was also investigated. Methylation levels at 165 CpG sites had Pearson’s correlation values of 0.9 or greater with sex (Table S3). Of these 165 sites, 160 were located on the X chromosome in humans (Table S3), one was located on the Y chromosome in humans, two were located on autosomal chromosomes in humans, and two did not have known coordinates in the human genome. The two autosomal sites included probe cg26452915 (Chr20:58911021, annotated as GNAS) and cg25449272 (Chr15:56654033, annotated as ZNF280D). A logistic model (*p*_*female*_ = 1/1+*e*^−(0.6717 – 1.1579*β-value)^) generated using a LASSO logistic regression machine learning method implemented in *cv.glmnet()* selected a single CpG: probe cg15451847, with a Pearson’s correlation of r = −0.999. The model predicted sex in the calibration samples (after thresholding the predicted probability > 0.5 indicating a female and outputs < 0.5 indicating a male). Probe cg15451847 corresponds to a site on the Y chromosome in humans (ChrY:19715996, annotated as KDM5D), indicating that its utility in predicting sex does not come from sexually dimorphic methylation patterns, but rather from the detection of a Y chromosome.

### Age-correlated CpG sites and the beluga epigenetic clock

The majority of CpG sites showed correlations with age of r < 0.5 (n = 28,232), and only 1.9% of CpG sites had a correlation of r > 0.5 (n = 551) (Table 1). No single site had a correlation coefficient larger than 0.9 but 21 sites exhibited correlations of between 0.8 and 0.9. The majority of CpGs assayed on the methylation array have negative correlations with age, and nearly 100% of the strongest correlations (> 0.7) were negative (Table 1).

**Table 1.** The frequency of absolute values of Pearson’s correlation coefficients (r) for the relationship between methylation (beta values) at 28,875 CpG sites and chronological age based on teeth growth layers of the 67 calibration samples, with age in 0.1 bins (each range is inclusive of the lower bound and exclusive of the upper bound). The right column shows the percent of the CpGs in each bin that have a negative correlation.

| Absolute value Pearson's $r$ | Number of CpG sites | % with negative correlation |
| --- | --- | --- |
| 0.0 - 0.1 | 14,133 | 56.5% |
| 0.1 - 0.2 | 8,663 | 59.9% |
| 0.2 - 0.3 | 3,887 | 58.1% |
| 0.3 - 0.4 | 1,200 | 67.1% |
| 0.4 - 0.5 | 427 | 78.9% |
| 0.5 - 0.6 | 253 | 87.7% |
| 0.6 - 0.7 | 173 | 96.0% |
| 0.7 - 0.8 | 115 | 99.1% |
| 0.8 - 0.9 | 24 | 100% |
| 0.9 - 1.0 | 0 | NA |
| Total | 28,875 | 59.2% |

Using elastic net regularization with values of alpha between 0.1 and 1 at 0.1 intervals, *cv.glmnet()* yielded models of varying sizes using between 20 and 134 CpGs with R^2^ values ranging from 0.923 to 0.942 (Table S4). The final model selected as the beluga epigenetic clock (alpha = 0.9) uses 23 CpG sites to generate age predictions, meaning the clock model has 24 terms, including the y-intercept (Table 2). The information in Table 2 comprises the multiple linear regression model and is all the information needed to calculate age for new samples using data from the custom methylation array. We selected this specific model to optimize for median age error, R^2^, and the y-intercept to reduce the tendency to overestimate the age of young whales. A linear regression of ages calculated with this model versus the tooth ages for each calibration sample resulted in a training set estimate of R^2^ = 0.92 (Figure 2A; other stats in Table 3). However, this training set estimate of the predictive accuracy is biased. To arrive at an estimate of the accuracy that is less biased by the nature of the training data, we employed leave one out cross-validation (LOOCV). The LOOCV run for an alpha of 0.9, which is intended to approximate the model’s performance on unknown data, had an R^2^ value of 0.74, a mean absolute age error of 3.65 years, and a median age error of 2.87 years (Fig 2B). The absolute age error for each sample in the LOOCV (the difference between the LOOCV predicted age and the estimated chronological age) showed a trend of underestimating the age of old whales while overestimating the age of young whales (regression slope = 0.65; regression y-intercept = 7.22).

**Table 2:** The CpG sites selected for the beluga epigenetic clock with associated model coefficients (to be multiplied by the CpG beta value), including the y-intercept. The CpG sites are referenced to the array probe names (Table S5). The Pearson’s correlation coefficient (r) for the methylation ratio with the tooth growth layer count is shown for each individual CpG site.

| Probe ID | Model Coefficient | CpG Correlation |
| --- | --- | --- |
| (Intercept) | 77.9708623 | NA |
| **cg00952468** | -4.525104457 | -0.7578 |
| **cg02534193** | -9.85801758 | -0.6457 |
| **cg02714609** | -14.10074358 | -0.8206 |
| **cg07279255** | 14.33543764 | 0.6956 |
| **cg07493173** | -0.175897809 | -0.7854 |
| **cg09622321** | -8.372865623 | -0.8686 |
| **cg12584622** | -6.025047791 | -0.8685 |
| **cg14043264** | -5.78974944 | -0.7817 |
| **cg14671961** | -5.341584389 | -0.6566 |
| **cg15809488** | -0.469547261 | -0.7487 |
| **cg15992086** | -0.379907718 | -0.6122 |
| **cg16678811** | -2.632026911 | -0.7887 |
| **cg17856858** | -1.525713524 | -0.8161 |
| **cg18629679** | -2.129559849 | -0.8387 |
| **cg21419180** | 30.73879442 | 0.4590 |
| **cg21420547** | 5.593128066 | 0.5444 |
| **cg22069272** | -5.967431136 | -0.8290 |
| **cg22416332** | -9.317426596 | -0.7402 |
| **cg25579908** | -19.26589983 | -0.7404 |
| **cg26286303** | -0.357946396 | -0.7184 |
| **cg26313355** | -1.762140707 | -0.7743 |
| **cg26899365** | -0.44830474 | -0.7246 |
| **cg27600712** | -0.104727313 | -0.8134 |

**Table 3.** Statistics for the beluga epigenetic clock model and leave one out cross-validation for alpha = 0.9: mean age error (mae), median age error (medae), r-squared for the regression, p-value for the regression, regression slope, and y-intercept for the regression.

| sex | mae | medae | R<sup>2</sup> | p-value | regression slope | y-intercept |
| --- | --- | --- | --- | --- | --- | --- |
| **Beluga epigenetic clock** |  |  |  |  |  |  |
| **all** | 2.34 | 1.97 | 0.92 | 3.50E-38 | 0.76 | 5.17 |
| **m** | 2.64 | 1.96 | 0.92 | 4.31E-20 | 0.73 | 5.89 |
| **f** | 1.99 | 2.06 | 0.94 | 6.33E-19 | 0.79 | 4.25 |
| **Leave one out cross-validation** |  |  |  |  |  |  |
| **all** | 3.65 | 2.87 | 0.74 | 1.14E-20 | 0.65 | 7.22 |
| **m** | 4.16 | 2.98 | 0.70 | 2.35E-10 | 0.63 | 8.01 |
| **f** | 3.06 | 2.56 | 0.81 | 7.58E-12 | 0.68 | 6.27 |

**Figure 2.**
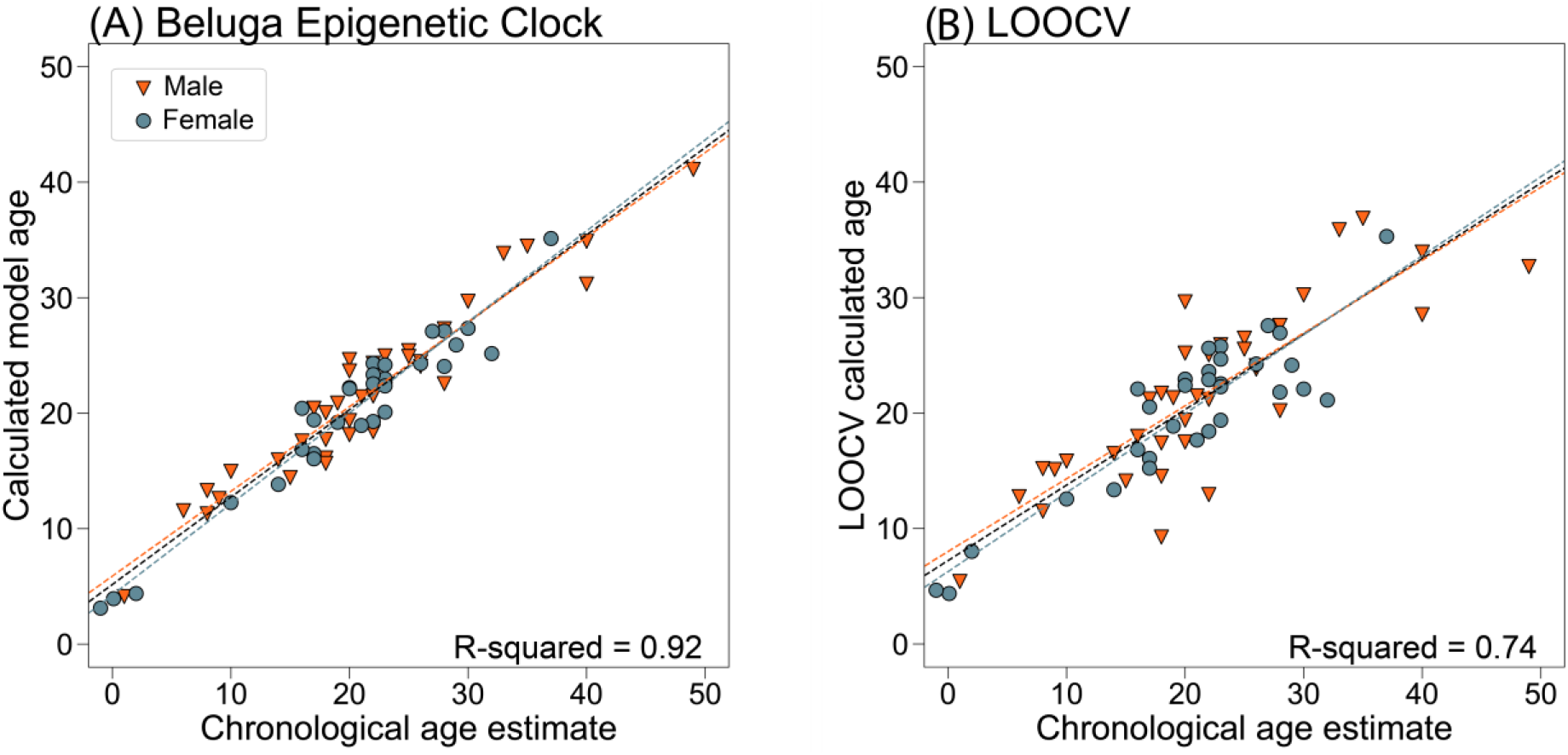
**(A)** Epigenetic ages calculated using the beluga epigenetic clock regressed against estimated chronological ages (based on GLG) for the calibration dataset. Data for males are represented by orange triangles and those for females are represented by grey-blue circles. Sex-specific regression lines as well as the overall regression line are shown (orange dashed line for males, grey/blue dashed line for females, black dashed line for overall regression). The training data showed an overall R^2^ = 0.92 (p = 3.50e^−38^). See Table 3 for all other statistics and sex-specific values. **(B)** Leave one out cross-validation (LOOCV) of the cv.glmnet() model parameters (alpha =0.7, lambada.min) with the same color scheme for males and females as panel A. Overall LOOCV R^2^ = 0.74 (p = 1.14e^−20^). See Table 3 for all other statistics and sex-specific values.

The R^2^ values for regressions between model predicted age and chronological age were consistently, but only slightly, higher for females than males in both the full clock model and LOOCV. None of the CpGs used in the final clock showed a sex correlation of 0.5 or greater.

### Genomic location and identity of clock CpG sites

The locations of all 23 CpG sites in the beluga genome were identified using BLAST and the NCBI genome viewer (Table S5). Annotations and gene ontology information for each CpG site revealed a wide range of gene identities and putative functions (Table S5). Of interest because of the role of epigenetics in gene regulation, four of the CpG sites fall in genes that have GO terms related to nuclear chromatin (cg00952468, cg15809488, cg21419180), promoter-specific chromatin binding (cg21419180), and chromatin remodeling (cg26899365). Fifteen of the 23 CpG sites were located within known genes that are annotated in the beluga genome; 18 of the sites are annotated within a gene in the human genome. Each annotation is unique, indicating that the CpGs are not linked within the same gene.

### Application of the beluga epigenetic clock to living beluga whales in the Cook Inlet

The methylation states of all 37,491 array CpGs were measured for genomic DNA from 38 skin tissue samples taken from living beluga whales. Data for the 23 clock CpG sites were used as input into the clock model yielding ages from approximately 11 to 27 years old, with potential range of +/− 2.9 years, using the LOOCV median age error of the calibration dataset (Fig 3; Table S6). The lower end of the estimated age distribution is consistent with the field practice of only sampling whales that are large juveniles or older. Additionally, epigenetic ages are in agreement with broad color categories that can be used to determine age classes of younger whales before they are entirely white (Wade, unpublished data; Figure S1).

**Figure 3.**
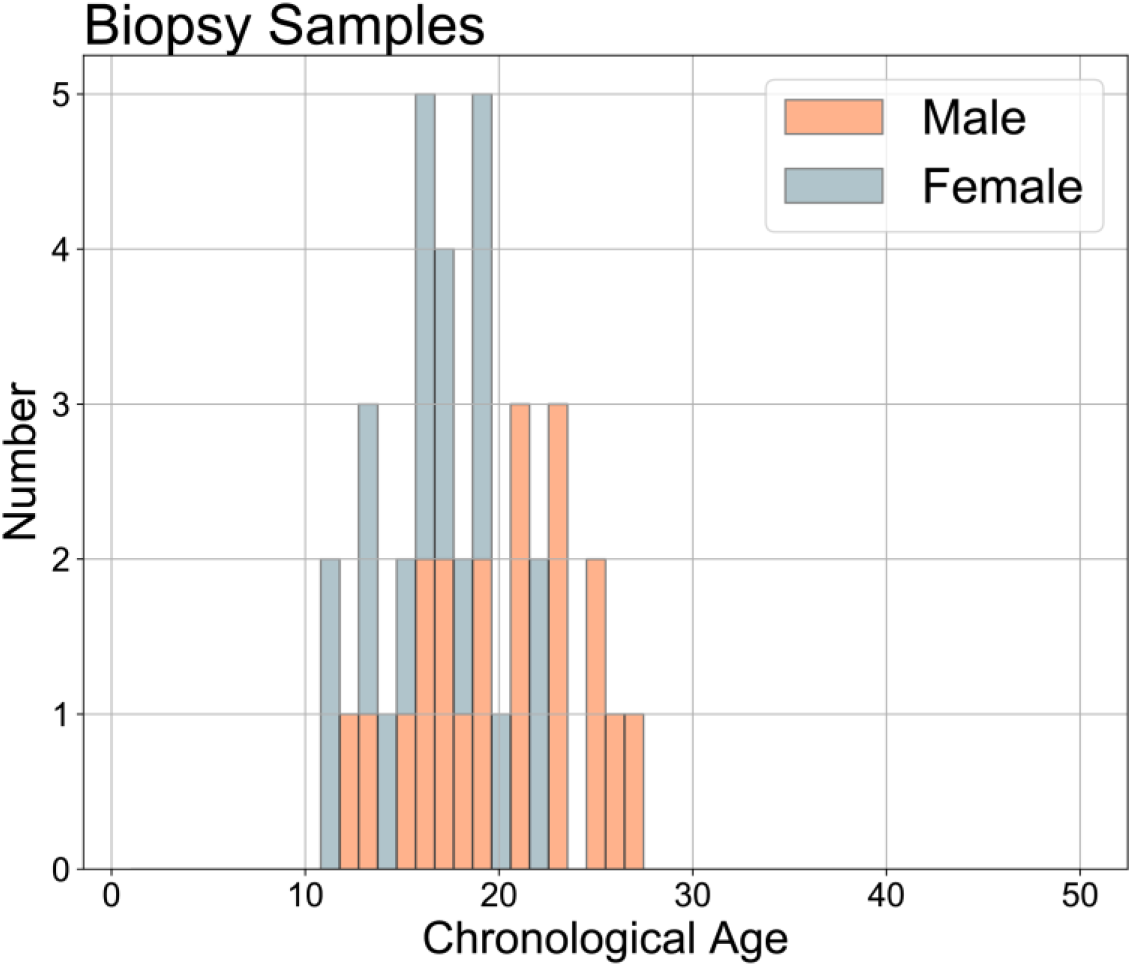
The estimated epigenetic ages for biopsy samples collected from living beluga whales between 2016 and 2018. Ages ranged from approximately 11 to 27. Note, this is not a representative age distribution of the populations due to bias in individuals biopsied during field seasons (e.g., younger whales not sampled with biopsy dart).

## Discussion

This study reports a robust epigenetic clock for beluga whales, enabling age estimation of living whales with just a small piece of skin tissue. The beluga epigenetic clock is based on 23 CpG sites that were selected from 37,491 CpG probes on a custom mammalian methylation array. Age estimation based on the multivariate age estimation model greatly outperforms age estimation based on a single CpG, which is consistent with what has been observed in other mammalian species including humans (Bocklandt, 2011; Hannum, 2013; Horvath, 2013; Thompson et al., 2018). The leave one out cross-validation (LOOCV) analysis suggests that this beluga epigenetic clock estimates age with a median absolute error of 2.9 years for samples of unknown age. Future independent test data are needed to fully validate the applicability of the clock to new datasets. Beluga whale longevity has been estimated to be 60 or 70 years (Suydam, 2009; Burns and Seaman, 1986), which would mean the clock approximates age within +/− 5% of the beluga lifespan.

The LOOCV showed a pattern of age underestimation for older whales and overestimation of younger whales, a pattern that is present but less pronounced in the training dataset. This pattern could partially be driven by data scarcity of older and younger whales in our calibration dataset, but it is also observed in other epigenetic clocks (e.g., Beal et al., 2019; Polanowski et al., 2014). Future research is needed to clarify the clock’s accuracy at the two ends of the age distribution. The young samples in our calibration dataset – one fetus and three calves – may have unique epigenetic changes occurring due to developmental processes or stress. Special considerations may be needed for fetuses and neonates from stranded mothers or still births.

It can be advantageous to carry out a non-linear transformation of age (e.g. a log transformation) to account for faster epigenetic changes occurring during development (Horvath, 2013; Hannum, 2013). Here, we did not carry out a non-linear age transformation because we found no evidence that it would improve the model fit in our data. This approach is consistent with other studies that directly regressed age on the CpGs (Polanowski et al., 2014; Thompson et al., 2018, 2017; Wright et al., 2018).

Future work that combines epigenetic methods presented here with research describing the relationship between morphometric features of beluga whale calves and age could offer a line of research that would improve clock performance for very young whales and fetuses (Robeck et al., 2015; Shelden et al., 2019). The development of alternative clock models could better describe variation in the rate of aging with life phase.

We found it important to prefilter the normalized CpG data based on detection p-value. Without any detection p-value filtering, we developed a clock with 59 CpGs (alpha = 0.5). While this clock led to an excellent predictive accuracy for chronological age as measured by R^2^ values (Table S2), we found that several of the underlying probes did not align to a CpG in the beluga genome. To alleviate concerns about overfitting, we built the final beluga clock (based on 23 CpGs) using only CpGs in a filtered dataset that required a significant detection p-value in at least 10 individuals.

Methylation data was also used to impute the sex of beluga whales. Many sex-associated sites are located on the X-chromosome in humans, but those that were not could be the focus of future study on sexual dimorphism in methylation patterns. Sex-based correlation analyses allowed us to assess the possibility that some CpGs were included in the clock due to sex-based patterns instead of age alone. This was not the case: none of the 23 CpGs in the clock had a Pearson’s correlation of more than 0.5 with sex. Furthermore, because there was no substantial sex-based difference in clock performance, our results support the use of this epigenetic clock for both sexes. While sexual dimorphism in morphology and behavior can be observed in beluga whales (Hauser et al., 2017), we found no indication that sex-specific clocks are required for estimation of chronological age. Furthermore, using one set of CpGs for both sexes will enable the development of an accessible age and sex assay in the lab by sequencing just 24 genomic regions (the 23 clock CpGs and the one sex-predictor CpG).

The sequencing of the beluga whale genome (Jones et al., 2017) increased the capacity for molecular research on this non-model species, enabling us to identify and map the 23 CpG sites in the beluga epigenetic clock. The 23 sites in the beluga epigenetic clock are found in genes related to critical biological activity like transcription, metabolism, cell membranes, etc. The function and mechanistic relationship of these genes with age is an open question. Clock development using an array, instead of targeted candidate genes, allows for comparison of important clock sites across mammal species without predisposing researchers to use just a handful of genes that have shown some relationship in the past.

The beluga epigenetic clock was successfully applied to skin tissue samples collected with a biopsy dart from living whales in Cook Inlet, Alaska. We present data from 38 skin tissue samples, but photo ID evaluation after the field season has indicated that three of these samples may be from the same individual (MML_RA180909_B01, MML_RA180910_B04, and MML_RA180912_B02). The estimated epigenetic ages for those three potential repeat samples are 26, 28, and 26; and the samples are all male, so the results perhaps support the photo identification results that this may be a recaptured whale. Genotyping is the best method to ultimately confirm. Age from biopsy samples will be useful in contributing to many different studies related to the conservation and management of beluga whales. With further development, it may be possible to partition CpG sites that correlate with chronological age from those that reflect biological age. Whereas ‘chronological age’ is important for demographics parameters, ‘biological age’ could be used to investigate the numerous physiological changes associated with aging (De Paoli-Iseppi et al., 2017; Horvath and Raj, 2018). In some populations or individuals, this biological aging is accelerated due to stress and exposure to environmental contaminants (a concept known as accelerated epigenetic aging). Future studies should explore whether epigenetic age acceleration can be observed in different whale populations, potentially reflecting genetic differences or various stress factors.

Research that compares epigenetic aging of Cook Inlet beluga whales with other populations will inform the applicability of this epigenetic clock to circumpolar populations of beluga whales. Data from other populations of beluga whales could also improve the accuracy of the beluga epigenetic clock by increasing sample size (the most accurate human clocks were trained on thousands of samples, e.g. Horvath, 2013), and help to mitigate error from chronological age estimates based on tooth growth layer groups. Tooth aging is subject to unknown error: beluga teeth wear with age at a rate that has not been quantified and is possibly individual-specific (Vos et al., 2019). Our analysis critically relies on the assumption that GLG patterns are well calibrated when it comes to estimating the chronological age of beluga whales. Beyond other beluga populations, this research also facilitates phylogenic comparisons of epigenetic clock CpGs with other cetacean species. Using a methylation array and machine learning to develop clocks will enable interspecific comparisons of age-relevant methylation patterns, potentially improving our understanding of the evolutionary function of age-correlated methylation.

## Acknowledgments

Funding was provided by the North Pacific Research Board (project #1723) and the Cooperative Institute for Marine Resource Studies (project #NB293T). SH and the generation of methylation data were supported by the Paul G. Allen Frontiers Group. We thank Debbie Steel for invaluable assistance in the Cetacean Genomics and Conservation Laboratory and Joe DeYoung in the University of California, Neuroscience Genomics Core. We thank the subsistence hunters in the Cook Inlet who provided samples of hunted whales for archiving. The use of both archived tissue samples (collected through the Alaska Marine Mammal Stranding Network under NMFS scientific research permits #932-1905/MA-009526) and modern biopsy samples (collected under NMFS Permit #14245-04) would not be possible without extensive field support from numerous scientists and organizations. NOAA’s Alaska Fisheries Science Center, Alaska Regional Office, Southwest Fisheries Science Center (Kelly Robertson), Northwest Fisheries Science Center (Kim Parsons), Group for Research and Education on Marine Mammals (Robert Michaud, Michel Moisan), Alaska SeaLife Center, Alaska Veterinary Pathology Services (Kathy Burek-Huntington), Joint Base Elmendorf Richardson (Christopher Garner), and the Cook Inlet Beluga Whale Photo-ID Project (Tamara McGuire) provided personnel, expertise, and/or data. The views implied or expressed here are those of the authors and do not necessarily represent the views of the National Marine Fisheries Service, NOAA. Reference in this document to trade names does not imply endorsement by the National Marine Fisheries Service, NOAA.

## Data Availability and Archiving Statement

All analysis scripts can be accessed on the Author’s GitHub at https://github.com/ekbors. Raw iDAT files are available at [embargoed until publication]. The manifest for the Horvath Methylation Array is available at Gene Expression Omnibus (GPL28271: Illumina HorvathMammalianMethylChip40 BeadChip). Processed data have also been archived in Axiom (link embargoed until publication).

## Supplemental Information

**Table S1:** This file contains detailed individual information for samples included in the clock calibration dataset: Southwest Fisheries Science Center (SWFSC) ID, Sample ID (includes extraction number when relevant), Sex, Age, Tooth Cap (presence of neonatal tooth cap), Length, Sample Location, and Sample Date.

**Table S2.** This file contains SeSAMe normalization detection p-values, the number of probes retained, and information on clock composition under various filtering scenarios. Data were generated for all alpha values, but for comparison to our published clock, here we share the clock data for alpha = 0.5 (default) and alpha = 0.9 (the final clock alpha).

**Table S3.** This file contains a list of the probe IDs with high correlations with sex, the Pearson’s correlation value (r), and the location of the probe in the human genome (sex-linked or not). All CpGs with a correlation > 0.5 are included in the sheet but annotations are only provided for sites with a correlation > 0.9.

**Table S4.** This file contains elastic net regularization clock data and performance statistics for all alpha values tested from 0.1 to 1 for the final calibration dataset clock models and LOOCV, with p-value filtering.

**Table S5.** This file contains the genome coordinates, gene annotations, and GO terms for the 23 clock CpGs.

**Table S6.** This file contains the ages of whales biopsied in 2016, 2017, and 2018 included in this study. Field ID information corresponds to the NOAA ID for each whale sampled.

**Figure S1.** This file contains field photographs of the youngest and oldest biopsied whales based on the epigenetic age estimation.

